# LysoTracker Deep Red exhibits photoconversion in multi-color structured illumination microscopy

**DOI:** 10.1101/2023.10.29.564502

**Authors:** Ida Sundvor Opstad, Kenneth Bowitz Larsen, Åsa Birna Birgisdottir, Krishna Agarwal

**Affiliations:** Department of Physics and Technology, UiT The Arctic University of Norway, NO-9037 Tromsø, Norway; Department of Medical Biology, UiT The Arctic University of Norway, NO-9037 Tromsø, Norway; Department of Clinical Medicine, UiT The Arctic University of Norway, NO-9037 Tromsø, Norway; Division of Cardiothoracic and Respiratory Medicine, University Hospital of North Norway, NO-9037 Tromsø, Norway

**Keywords:** lysotracker deep red, photoconversion, structured illumination microscopy

## Abstract

In new chemical environments or for untested combinations of illumination, unexpected changes to the fluorescent labels’ photophysical properties, such as photoconversion, can occur. This letter reports on photoblueing of a common cellular probe, LysoTracker Deep Red, in multi-color super-resolution structured illumination microscopy. The dye was found to exhibit a blue shift of its fluorescence spectrum as a step on the photobleaching pathway during such imaging. The observed photoblueing (emission spectrum shifted to lower wavelengths) of this probe is an important finding as many cellular assays rely on the spectral separation of this and such fluorescent cellular markers. No spectral shift of LysoTracker Deep Red was observed for longterm imaging using diffraction limited microscopy requiring significantly lower light dose. We expect that the knowledge of occurrence of photoconversion during super-resolved imaging using this popular dye will help researchers design better imaging experiments and avoid potentially erroneous interpretations in multi-color microscopy.

## 1. INTRODUCTION

The possibility of specific labelling and visualization of subcellular molecules and compartments have made super-resolution microscopy an invaluable tool in biological research [1]. The sought image specificity can however be obstructed by several factors like unspecific labelling, spectral overlap in multi-color experiments, or autofluorescence of cells and tissues [2]. Another factor that can misguide biological inferences is the undesired photoconversion of fluorescent labels. Photoconversion involves a spectral shift of the initial fluorophore induced by light exposure. The local environment and fluorophore structure have a strong impact on both photobleaching and photoconversion. Photoconversion can for some fluorophores be a step in the photobleaching pathway [3]. Photoconversion has been studied in the context of single-molecule localization microscopy during or after intense laser-exposure [4]. Further, the photoblueing of the dye Lysotracker Red (a shift from red to green) has been reported after illumination with an epifluorescence light source [5].

In this work, we report on photoconversion of the commonly used cellular probe LysoTracker Deep Red (LTDR) for the volumetric super-resolution imaging technique three-dimensional structured illumination microscopy (3DSIM) [6]. 3DSIM is considered relatively low intensity and live-cell friendly compared to other super-resolution microscopy techniques [1]. Still, the repeated light exposure required for multi-color, volumetric and time-lapse imaging has been seen to cause both rapid photobleaching and phototoxic effects in living cells [7]. Lastly, the photoblueing of LTDR in long-term diffraction limited (conventional) microscopy requiring significantly less light exposure is evaluated. Both the conventional microscopy and 3DSIM data used for this work are published as open datasets [7, 8].

## 2. RESULTS

H9c2 cardiomyoblasts were stably transfected to express the double tag mCherry-EGFP on mitochondria and further incubated with LTDR for the visualization of acidic vesicles (primarily lysosomes). The double tag can be used to study the degradation of mitochondria in acidic vesicles, as the EGFP will be quickly quenched in an acidic environment while the mCherry red fluorescent protein has high acid stability [8]. Sequential 3DSIM imaging of the three channels was conducted in time-lapse mode with as low as possible illumination intensities while still achieving satisfactory SIM reconstruction quality. The 3DSIM images were investigated for possible mitochondrial uptake in the acidic compartments. All bright LTDR-positive vesicles investigated were found to quickly become red (instead of far/deep red) during time-lapse imaging. This could indicate very rapid uptake of mitochondrial fragments into lysosomes. However, the rate of the red signal increase in lysosomes was independent of the distance from mitochondria. Thus, the most plausible explanation for this observation is photoblueing of the LTDR dye. This observation is illustrated in **Figure 1** for a cell with many bright acidic compartments (bleach-corrected image sequence). One of the acidic vesicles is highlighted in the top right panel of Figure 1 at t1. While the mCherry channel initially does not show the vesicular geometry witnessed in the LTDR image, within 9 frames, we see the geometry being replicated in the mCherry/RFP channel as well. The same result is seen across many lysosomal structures within 9 time points. The bottom row shows a magnified view of the same vesicle, but for the raw data to avoid any possible effects of 3DSIM reconstruction artifacts especially in the photobleached images. The raw data intensity of this vesicle was measured to be approximately constant over time in the mCherry/RFP channel, while strongly decreasing for the LTDR/Cy5 channel (graph at lower right in Figure 1). The mCherry-labelled mitochondria, however, are strongly photobleaching, which can be seen from the non-bleach-corrected image sequence provided in **Supplementary Movie 1**. Note also the vesicles emerging in the RFP channel (in the movie shown in yellow) as the mitochondria fade away.

**Figure 1:**
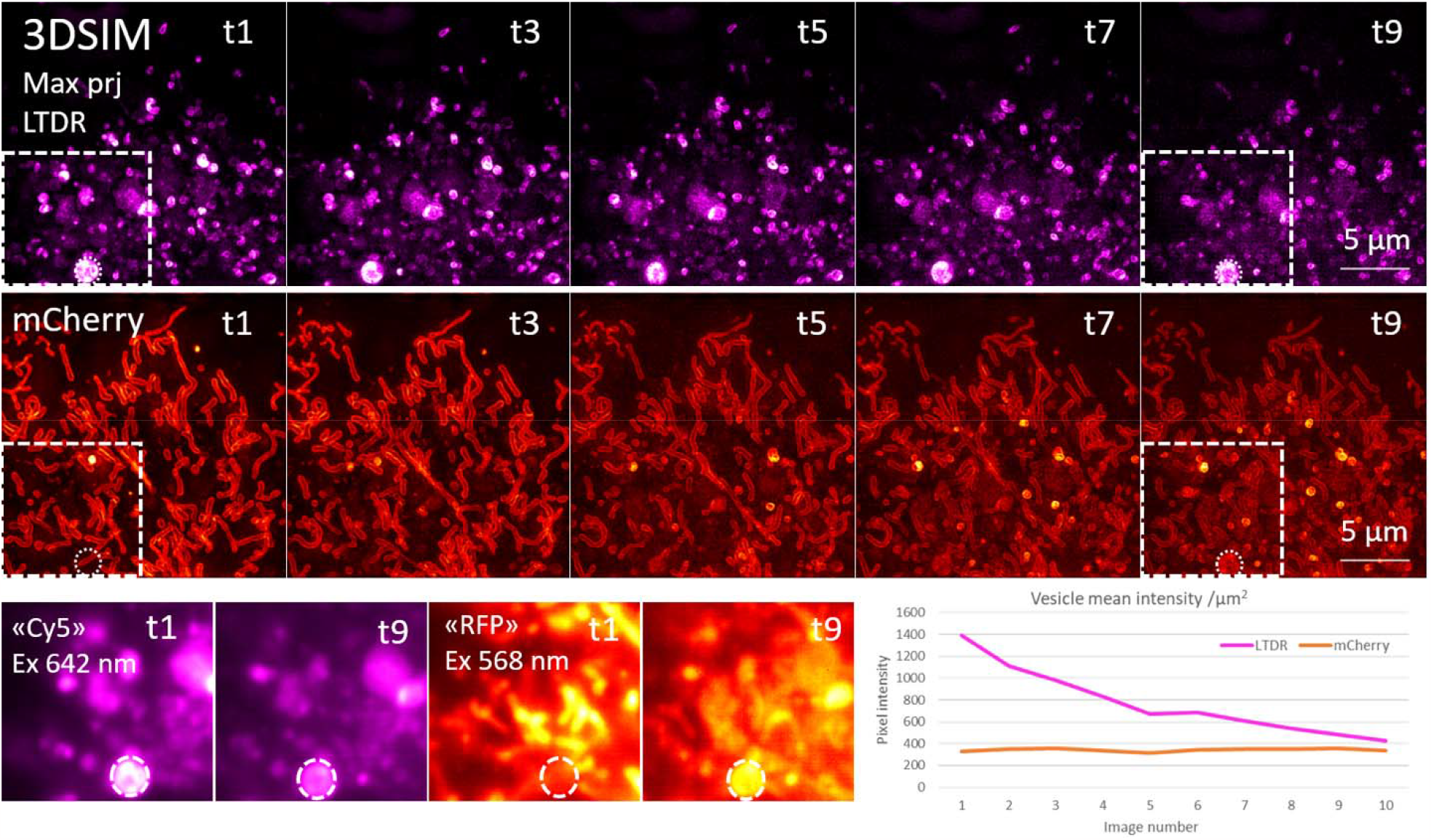
Photoblueing of LysoTracker Deep Red from the “Cy5” to “RFP” channel during multi-color 3DSIM acquisition. During the image sequence (t1 to t9 indicate time point), the bright image features from the Cy5 channel (acidic vesicles) are seen appearing in the RFP channel blended with signal from mCherry-labelled mitochondria. The cells were also exposed by 3DSIM imaging by a 488 nm laser during this sequence, which might affect the photoconversion. The two top rows show bleach corrected and maximum intensity z-projections of the reconstructed 3DSIM super-resolved images. Both channels are photobleaching, but the relative intensity of the RFP channel is increasing inside lysosomes. The bottom row images show mean intensity z-projection of the raw 3DSIM data from which the intensity measurements on the bottom right are taken. The graph illustrates the change in vesicle intensity over a sequence of 10 multi-color 3DSIM volumes (excitation: 568 nm and 642 nm). While the intensity in the far red “Cy5” channel is strongly decreasing, the red “RFP” channel is kept at a stable level due to photoconversion of LTDR from “Cy5” to “RFP”.

We further investigated if conventional fluorescence microscopy with considerably less light exposure than 3DSIM would show a similar effect on LTDR during a multi-color timelapse. An example from one such volumetric and multi-color time sequence is displayed in **Figure 2** for time point 1, 30 and 60. Different from the 3DSIM images in Figure 1, this time-sequence is not bleach corrected as the photobleaching is very slow. For this timelapse, no obvious photoconversion of LTDR to the RFP channel is visible.

**Figure 2:**
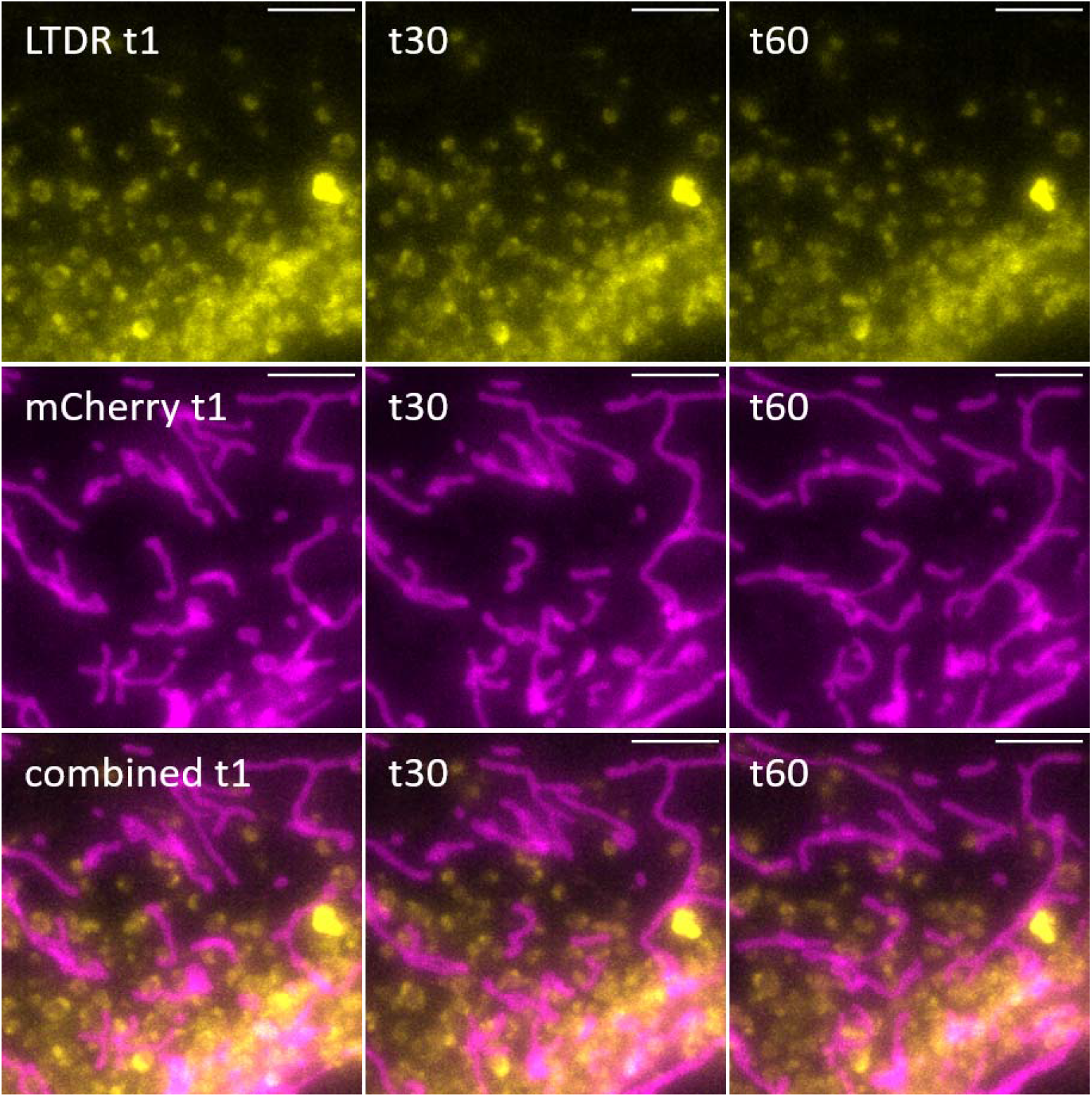
Three time points from a conventional fluorescence microscopy timelapse (30 volumetric timepoints apart) of LTDR and mitochondria (mCherry). The images are maximum intensity z-projections. In addition to the “Cy5” and “RPF” channels, as for 3DSIM, also “GFP” was excited and imaged (GFP not shown). For the conventional time-lapse, no photoconversion of LTDR is visible. The scale bars are 5 μm.

Considering the illumination modules and experimental settings for the 3DSIM and the conventional imaging, the illumination power used by the two techniques are comparable in the range of 0.55 mW – 5 mW for 3DSIM and 2.2 mW – 13 mW for conventional imaging for the three channels, both techniques with 5 ms to 10 ms camera exposure time. More details about the illumination modules are provided in the methods section. Although the illumination power on the sample as a whole is similar for the two techniques, the local intensities (power per area) experienced by the individual cells or molecules are likely very different, as 3DSIM uses coherent and polarized illumination focused onto the sample for the repeated generation of illumination fringes covering only 2-3 cells at the time (the illumination fringes can be seen to cover a circle of about 82 μm diameter when using the full camera field-of-view). The conventional illumination on the other hand, is not coherent and cannot (and need not) be focused as tightly as for 3DSIM (the illumination area is much larger than the camera field-of-view).

Another significant difference, which is also easier to compare, is the number of images acquired per volumetric time-point. For both methods, 2 μm deep z-stacks were acquired, for conventional microscopy corresponding to 9 z-planes and for 3DSIM 17 z-planes (the higher z-resolution for 3DSIM requires higher z-sampling rate). Additionally, for each super-resolved image plane, this 3DSIM acquires 15 images (3 angles and 5 phase shifts of the excitation interference pattern). Per volume, this then multiplies to 255 images per color channel for 3DSIM, but only 9 images for conventional fluorescence. During the 10 time-points 7 s timelapse shown in Figure 1, images were acquired continuously (3 x 255), making this the fastest time-lapse possible for this multicolor acquisition. In comparison, the conventional imaging underlying Figure 2 was a 5 s timelapse and allowed for some illumination pause between the acquisitions.

To summarize, despite similar powers provided by the two different illumination modules, the observed photoconversion for 3DSIM but not for conventional imaging, can be attributed to the more focused power (higher local intensities) by 3DSIM and/or the 28 times more light exposures per volume for 3DSIM acquisition. Differences in how a polarized, coherent light source compared to an incoherent LED illumination affect the photoconversion of the fluorescent molecules might also contribute to the observed differences between 3DSIM and conventional fluorescence microscopy.

## 3 CONCLUSION

We have in this letter given a first report of photoconversion of the cellular probe LysoTracker Deep Red in multi-color 3DSIM video microscopy. During three channel imaging (excitation using 488, 568, and 642 nm), the bright vesicular structures in the Cy5 channel were observed to blead into the RFP channel perturbing the intended analysis of mitochondria degradation. This 3DSIM image sequence was rapidly photobleaching and had to be bleach-corrected for visualization.

We further investigated if the same mitochondria-lysosome assay could be utilized in conventional fluorescence microscopy with significantly less light exposure than 3DSIM. For these 60-time point-long multi-color volumetric videos, the photobleaching was negligible and we could not find any signs of a similar photoblueing as for the 3DSIM data.

We further investigated the differences in illumination between the two imaging modes. The 3DSIM acquisition was found to use less power onto the sample as a whole per camera exposure but acquires 28.3 times more images per image volume compared to conventional widefield microscopy in addition to using a focused laser beam for sample illumination, increasing the local intensity experienced by individual cells and dye molecules.

In conclusion, the photoblueing of LTDR appears to be a step in the photobleaching pathway of LysoTracker Deep Red but the dye still appears safe to use in multi-color assays as long as the photobleaching is negligible. Finally, we encourage everyone to critically evaluate their assays and multi-color image data for similar photoconversion effects that might hamper biological interpretations.

## 4 EXPERIMENTAL SECTION

### 4.1 Sample preparation

The rat cardiomyoblast cell-line H9c2 (cells derived from embryonic heart tissue; Sigma Aldrich) was genetically modified using retrovirus to achieve a stable expression of tandem tagged (mCherry-EGFP) mitochondrial outer membrane protein 25 (OMP25)-transmembrane domain (TM). A uniform expression of fluorescence intensity in the cells was achieved through flow cytometry sorting. The cells were incubated with LysoTracker Deep Red (LTDR, Cat nr L12492, Thermo Fisher) for the visualization of acidic vesicles (100 nM for 40 min for 3DSIM, 50 nM for 30 min for widefield. Detailed sample and experimental protocols are available with the published datasets [7, 8]. The 3DSIM data is further described in [9].

### 4.2 Experimental data

The data used for this article is for 3DSIM from [10], and for conventional fluorescence from [11]. For Figure 1 (3DSIM) the file 20210511_H9c2-dTag_GLU_LTDR100nm-40m_1520_sim256_005 was used. For Figure 2, the file 20210330_H9c2_mito-mChGFP_LTRDR50nM_1522_dvt-5sTL_005 was used.

### 4.3 Microscope

3DSIM and conventional fluorescence imaging was conducted using an OMX Blaze v4 3DSIM system (in conventional or 3DSIM mode).

#### 3 DSIM mode

3DSIM excitation lasers: 488 nm (100 mW with 10% transmission, 5 ms exposure), 568 nm (100 mW with 10% transmission, 10 ms exposure), and 642 nm (110 mW with 1% transmission, 5 ms exposure). At best 50 % of the transmitted laser power is expected to reach the sample after optical fibers and illumination engineering. For each color and each plane, 15 raw SIM images were acquired (three angles and 5 phases). The three-color volumetric images have a thickness/depth of 2 μm, corresponding to 17 z-planes 125 nm apart. The time between each multi-color volume was about 7 s.

#### Conventional/widefield mode

Illumination source: InsightSSI, a solid state illumination system. Excitation light: FITC-GFP (max. 110 mW, 5% transmission, 10 ms exposure), A568-mCherry (max. 125 mW, 10% transmission, 10 ms exposure), Cy5 (max. 45 mW, 5% transmission, 5 ms exposure). The image volume thickness was 2 μm, corresponding to 9 z-planes 250 nm apart. The time between each multi-color volume was 5 s.

### 4.5 Image analysis

The data was visualized and analyzed using Fiji [12].

## ACKNOWLEDGMENTS

KA acknowledges ERC Starting grant fund 804233 (3D-nanoMoprh) and Research Council of Norway’s FRIPRO Young fund 288082 (nanoRIP). ISO and KA acknowledge funds from UiT’s Tematiske Satsinger project with Cristin ID 2061348 (VirtualStain). ÅBB and KBL acknowledge funding from Helse Nord RHF [HNF1449-19].

## Abbreviations

Lysotracker Deep Red: LTDR
3DSIM: three-dimensional structured illumination microscopy

